# Similar patterns of background mortality across Europe are mostly driven by drought in European beech and a combination of drought and competition in Scots pine

**DOI:** 10.1101/551820

**Authors:** Juliette Archambeau, Paloma Ruiz Benito, Sophia Ratcliffe, Thibaut Fréjaville, Alexandre Changenet, Jose M. Muñoz Castañeda, Aleksi Lehtonen, Jonas Dahlgren, Miguel A. Zavala, Marta Benito Garzón

**Author notes:** Corresponding author: Juliette Archambeau.

## Abstract

**Aim:** Background tree mortality is a complex demographic process that affects forest structure and long-term dynamics. We aimed to test how drought intensity interacts with interspecific and intraspecific competition (or facilitation) in shaping individual mortality patterns across tree species ranges.

**Location:** European latitudinal gradient (Spain to Finland).

**Time period:** 1985 – 2014.

**Major taxa studied:** Scots pine (*Pinus sylvestris* L.) and European beech (*Fagus sylvatica* L.).

**Methods:** We performed logistic regression models based on individual tree mortality recorded in five European National Forest Inventories. We computed the relative importance of climatic drought intensity, basal area of conspecific and heterospecific trees (proxy of indirect intra- and interspecific competition or facilitation) and the effects of their interactions on mortality along the latitudinal gradient of both species range.

**Results:** Increase in drought intensity over the study period was associated with higher mortality rates in both species. Drought was the most important driver of beech mortality at almost all latitudes while Scots pine mortality was mainly driven by basal area. High conspecific basal area was associated with high mortality rates in both species while high heterospecific basal area was correlated with mortality rates that were high in Scots pine but low in beech.

**Main conclusions:** Beech mortality was directly affected by drought while Scots pine mortality was indirectly affected by drought through interactions with basal area. Despite their different sensitivity to drought and basal area, the highest predicted mortality rates for both species were at the ecotone between Mediterranean and cool temperate biomes, which can be explained by the combined effect of drought and competition. In the context of global warming, which is expected to be particularly strong in the Mediterranean biome, our results suggest that populations at the southern limit of species ranges may experience increased mortality rates in the near future.

**BIOSKETCH:** The authors’ research is focused on functional trait ecology and global change, with special attention to mortality and demography processes. The authors use modelling multidisciplinary approaches to understand complex processes in ecology on a large geographical scale.

## 1 INTRODUCTION

Tree mortality plays a major role in shaping forest dynamics, structure and composition (Franklin *et al.*, 1987; Ruiz-Benito *et al.*, 2017a), species range shifts (Benito Garzón *et al.*, 2013), ecosystem functioning and services (Millar & Stephenson, 2015), carbon fluxes and feedback to the global climate system (Sitch *et al.*, 2008). Therefore, understanding and predicting tree mortality is a key challenge in ecology, particularly in a changing climate.

Global change is exacerbating drought-induced tree mortality (Allen *et al.*, 2015). Recent forest die-off events have occurred in all major biomes and on every wooded continent (Allen *et al.*, 2015) and background tree mortality also appears to have increased in North America (Mantgem *et al.*, 2009; Hember *et al.*, 2017) and in Spain (Carnicer *et al.*, 2011). Yet, an empirical quantification of background tree mortality at continental scale is missing and whether or not forest mortality follows an increasing global trend that will keep rising under global change remains unclear (Hartmann *et al.*, 2018). Moreover, mortality is a major process which delimits species range (Gaston, 2009), notably at the driest edge of their distribution (Benito Garzón *et al.*, 2013). Therefore, large scale studies that capture the entire species distribution are essential to determine how climate change induced mortality might affect species distribution.

Understanding and predicting background tree mortality patterns at large scales remains challenging for several reasons. First of all, mortality is a stochastic phenomenon (Franklin *et al.*, 1987), which is therefore difficult to predict. Secondly, it is often the result of a complex and gradual process with multiple interacting drivers (Manion, 1981), that act at different spatial and temporal scales (Dietze & Moorcroft, 2011). Thirdly, there may be a lag time between episodic stressful conditions and tree mortality responses (Jump *et al.*, 2017). Lastly, background tree mortality rates are difficult to estimate due to the small sample size of dying trees in local studies, while large samples are needed to understand mortality patterns.

In European forests, background tree mortality is strongly driven by climate variability (Neumann *et al.*, 2017). Among the climatic factors affecting tree mortality, drought plays a major role (Benito Garzón *et al.*, 2013; Ruiz-Benito *et al.*, 2013; Allen *et al.*, 2015) and particularly affects populations at the driest edge of species distributions (Benito Garzón *et al.*, 2018). Among the biotic factors, competition for limited resources may be an important cause of tree mortality and may also interact with climate, notably through a higher increase in mortality rates in areas that are both dry and dense (Ruiz-Benito *et al.*, 2013; Vilà-Cabrera *et al.*, 2013; Young *et al.*, 2017). Moreover, tree mortality responses can differ widely depending on whether we consider intra- or inter-specific competition (Condés & del Río, 2015). However, how intra- and interspecific competition interact with climatic drought to shape range-wide mortality patterns remains unknown.

Tree mortality sensitivity to biotic and abiotic factors vary along species’ ecological strategies, from stress-tolerators to competitors and from angiosperms to gymnosperms (Choat *et al.*, 2012; Ruiz-Benito *et al.*, 2017a). European beech (*Fagus sylvatica* L.) and Scots pine (*Pinus sylvestris* L.) are two widely distributed European tree species with different life history strategies. Beech is a highly competitive, shade-tolerant and late-successional species, which is known to be sensitive to drought (San-Miguel-Ayanz *et al.*, 2016). Scots pine is considered as a weakly competitive, highly stress-tolerant and light demanding pioneer tree, which grows under both wet and dry conditions (San-Miguel-Ayanz *et al.*, 2016). Regional scale studies suggested that both species are being progressively replaced by other species in the southern part of their distribution (Vilà-Cabrera *et al.*, 2013; Galiano *et al.*, 2010) and in some inner Alpine valleys in the case of *P. sylvestris* (Rigling *et al.*, 2013).

Our main objective was to understand and predict range-wide patterns of background mortalities in Scots pine and European beech. To that end, we parameterised individual-level logistic regression models, as a function of climatic drought and tree basal area (used as a proxy of competition or facilitation), using records from five National Forest Inventories covering the entire European latitudinal gradient, from Spain to Finland. We hypothesised that (i) mortality in both species is influenced by climatic drought, basal area and their interaction but with a higher influence of basal area in the case of Scots pine; and (ii) that despite these differences in their sensitivity to drought and basal area, both species display similar spatial patterns of mortality across their ranges: high mortality in the south resulting from increasingly dry climates, especially in the Mediterranean biome.

## 2 METHODS

### 2.1 Forest inventory data

We used mortality data from five National Forest Inventories (NFIs) covering the entire European latitudinal gradient, from the Mediterranean to boreal biome. Data from four of the NFIs had been previously harmonised as part of FunDivEUROPE project (Spain, Germany, Sweden and Finland) and the French NFI was added to this study. In each NFI, trees were recorded in temporary or permanent plots depending on the country. Plots in the German, Finnish and Swedish inventories are gathered within clusters (see Appendix S1 for details of the survey design and sampling methods for each NFI). We selected plots in which at least one of our two target species (i.e. *F. sylvatica* or *P. sylvestris*) was recorded. These plots were classified into Mediterranean, cool temperate and boreal biomes (Fig. S1.1) and were unevenly distributed along the latitudinal gradient (Fig. S1.2). The final datasets contained 57,191 beech trees and 161,720 Scots pine trees in 10,150 plots and 16,669 plots, respectively. From those trees, 1,490 (2.6%) and 7,649 (4.7 %) were recorded as dead for beech and Scots pine, respectively.

As explanatory variables of tree mortality, we selected tree *DBH* (diameter at breast height) as *DBH* is known to influence individual tree mortality (Ruiz-Benito *et al.*, 2013). We additionally calculated three proxies of indirect competition between trees (or facilitation) (Fig. S2.2): basal area of neighbouring trees considering all tree species (i.e. *BAall*, m^2^ ha^−1^), basal area of neighbouring conspecifics (i.e. *BAintra*, m^2^ ha^−1^) and basal area of neighbouring heterospecifics (i.e. *BAinter*, m^2^ ha^−1^).

### 2.2 Drought-related variables

Climatic drought intensity over the study period (Fig. S2.1) was characterised by a water availability index: *WAI* = (*MAP*-*PET*) / *PET,* where *MAP* is the mean annual precipitation (mm) and *PET* the mean potential evapotranspiration (mm). For each plot, *PET* was extracted from the CRU v3.24.01 monthly gridded dataset at 0.5-degree resolution (Harris *et al.*, 2013) and *MAP* was calculated from a downscaled version of E-OBS at 1 km resolution (Moreno & Hasenauer, 2016). For each plot, *WAI* was averaged over the period between two years before the first survey date and the second survey date to include delayed effects of drought on mortality (Greenwood *et al.*, 2017).

Changes in climatic drought intensity over the study period (i.e. temporal variability of drought intensity) were described by the Standardized Precipitation Evapotranspiration Index (*SPEI,* Fig. S2.1; Vicente-Serrano *et al.*, 2009), obtained from a gridded dataset at 0.5-degree resolution (Beguería & Vicente Serrano, 2017). *SPEI* is a multi-scalar drought index whose variations have been shown to be highly correlated with tree response to climate (Greenwood *et al.*, 2017). Its calculation considers both *PET* and *MAP*, with *PET* derived from the Penman-Monteith equation. *SPEI* compares drought intensity during a long-term reference period (i.e. from 1901 to 2015) to that of a given period from 3 to 48 months. In our study, we selected a 12-month period to consider both current and previous year water shortage. *SPEI* is expressed as a standardised index relative to each site, with a standard deviation of 1, where negative values indicate more intense drought over the timescale considered compared to reference conditions. For each plot, we calculated mean *SPEI* (hereafter *SPEI*) over the period from two years before the first survey date to the second survey date.

### 2.3 Model description

We parameterised two species-specific models, where *P*_*i*_ is the annual probability of mortality for each individual tree *i*. We used a logistic regression model with a link *cloglog* to allow the sigmoidal curve of the mortality probability to be asymmetrical and deal with zero inflated distributions (Zuur *et al.*, 2009):

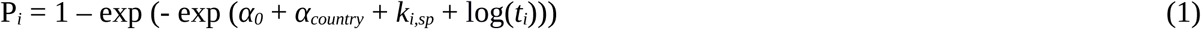

where *α*_*0*_ is an intercept term (set to zero); log(*t*_*i*_) is an offset variable that takes into account the survey interval length *t*_*i*_ (years) for each tree *i*; α_*country*_ is the random country intercept to include the sampling differences between each NFI and *k*_*i,sp*_ is a species-specific linear function that includes the relationship between the mortality of tree *i* of the species *sp* (i.e. *F. sylvatica* or *P. sylvestris*) and the explanatory fixed-effect variables. Although clusters and plots could be considered as a source of variation for each tree, we did not consider cluster and plot as random terms because most of the clusters contained only one plot and in many plots no trees died between the two survey dates. We used the function “glmer” of the “lme4” package to run the model described in equation 1 in R 3.3.3 (R Core Team 2017).

For both species, we explained mortality patterns using four fixed-effect predictors with low collinearity (i.e. Spearman correlation coefficient: r < 0.59, and Variance Inflection Factor: VIF < 2; Dormann et al., 2013), namely: *BAintra, BAinter, WAI* and *SPEI.* Conspecific and heterospecific basal area (i.e. *BAintra* and *BAinter*) were both included in the model as they can have different effects on tree mortality (Condés & del Río, 2015). To ensure a linear relationship between each explanatory variable and tree mortality, *BAinter, WAI* and *SPEI* were log transformed (see Appendix S3 for details).

Tree size (*DBH*) was included as a covariate in our model, as we were not directly interested in the importance of tree size on mortality. As we required a single parameter per predictor to estimate the relative importance of each predictor (see section 2.5), we calculated a non-linear variable from *DBH*: *DBHnl*_*sp*_ *= DBH + r*_*sp*_ *×* log(*DBH*) (see Appendix S3 for details).

To understand how tree mortality was affected by basal area and climatic drought, we included the main effect of each variable and first-order interaction terms between abiotic and biotic variables. Herewith, the function *k* from equation 1 took the form:

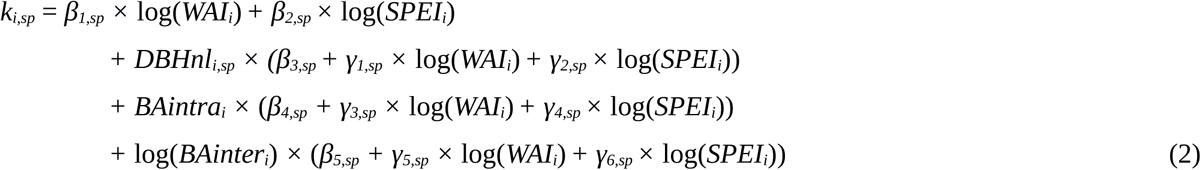

where *β*_*x*_ and *γ*_*x*_ are the estimated coefficients of the main and interaction effects, respectively (Table S3.1).

### 2.4 Model performance and evaluation

Binned residuals plots were used to ensure our final species-specific models were well-calibrated (Fig. S3.3-4). To evaluate the discrimination accuracy of our models, we computed the mean area under the curve (AUC) on 100 bootstrap samples among the predicted and observed values. AUC values of 0.6-0.7 show a fair discrimination accuracy, between 0.7 and 0.8 good and above 0.8 excellent (Hurst *et al.*, 2011). We used independent cross-validation to measure the generalisation power of the model, for which we used 75% of the data to fit the model and the remaining 25% to independently validate our predictions.

### 2.5 Relative importance of climatic drought and basal area on mortality

Following Ratcliffe *et al.*, (2016), we explored the relative importance of each predictor on individual tree mortality in relation to the other predictors by considering the predictors’ main effects and their interactions. For doing so, we first computed the absolute importance of each predictor using our model coefficients. For instance, to compute *A*_*BAintra,i*_ the absolute importance of *BAintra* on the probability of mortality of the tree *i*, we applied the following equation separately for each species:

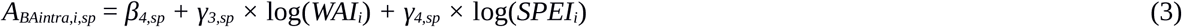

where *β*_*x*_ and *γ*_*x*_ are the estimated coefficients of the single predictors and their interaction effects respectively; *WAI*_*i*_ and *SPEI*_*i*_, are the plot values corresponding to these variables.

Secondly, the relative importance of each predictor was computed for each tree by dividing the absolute importance of the focal predictor by the maximum absolute importance between all predictors of the target tree. For instance, to estimate the relative importance of *BAintra* for the tree *i*, we calculated for each species: |*A*_*BAintra,i,sp*_| / max(|*A*_*BAintra,i,sp*_|, |*A*_*SPEI,i,sp*_|, |*A*_*WAI,i,sp*_|, |*A*_*BAinter,i,sp*_|); where *A*_*SPEI,i*_, *A*_*WAI,i*_ and *A*_*BAinter,i*_ are the absolute importance of *SPEI, WAI* and *BAinter* for tree *i*, respectively. For each tree *i*, the predictor that had the greatest influence on individual tree mortality probability had a relative importance of one.

## 3 RESULTS

### 3.1 Model performance and validation

Scots pine and beech models showed good agreement between observed and predicted values (AUC = 0.73 and 0.71, respectively). The Scots pine model performed well in predicting tree mortality probabilities across the European latitudinal gradient as predicted and observed values exhibited similar patterns (Fig. 1a). Nevertheless, caution is needed to interpret the results at the latitudinal gradient edges where Scots pine mortality rates were slightly underestimated. The beech model overestimated mortality rates in northern Germany and in Spain but accurately predicted mortality rates in the south of France and in the south of Sweden, at the north end of the gradient (Fig. 1b). Model and partial residual plots for each predictor showed no strong spatial patterns, thus supporting the validity of the models (Fig. S3.3-4).

**Figure 1.**
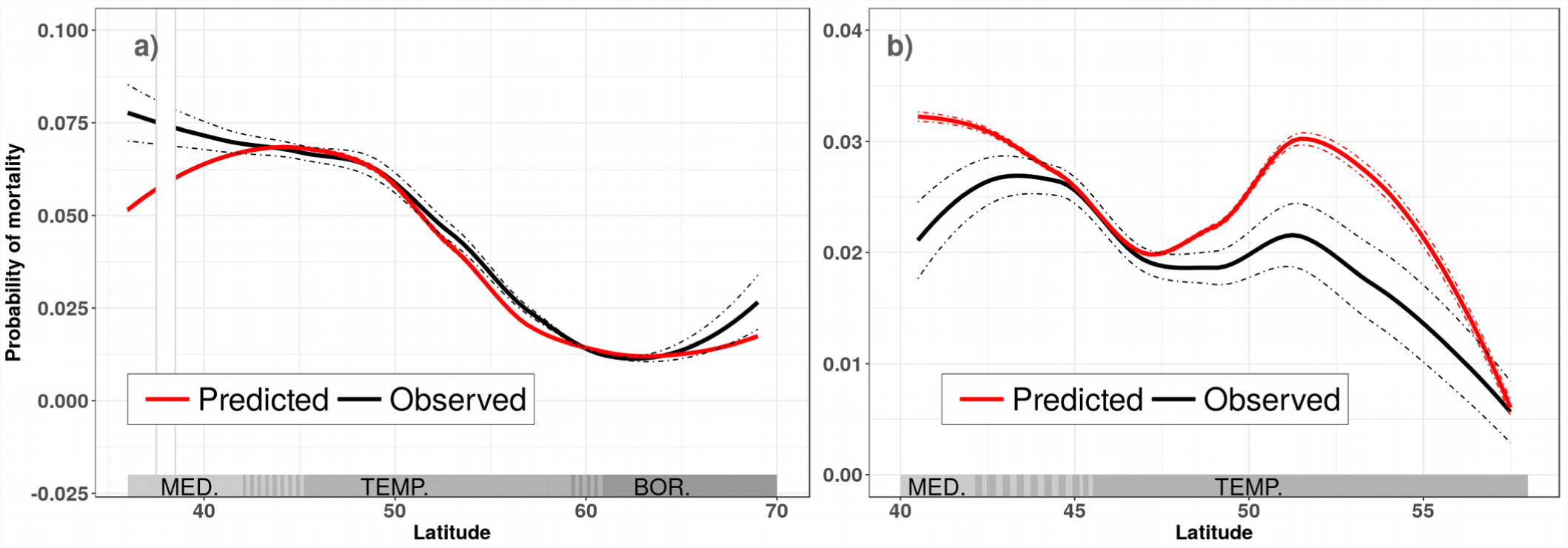
Predicted and observed individual probability of mortality along the latitudinal gradient covered by the NFIs plots a) for *P. sylvestris* and b) for *F. sylvatica*. Predicted values correspond to the predicted probability of mortality of each tree between the two survey dates. Predicted and observed values were clustered at 1° latitude for *P. sylvestris* (a) and at 0.5° resolution for *F. sylvatica* (b). A locally weighted regression was used to obtain the smooth solid lines (“loess” method of the geom_smooth function in “ggplot2” R package). Dotted lines indicate 95% confidence intervals. The acronyms MED., TEMP. and BOR. in grey bars refer to the Mediterranean, cool temperate and boreal biome, respectively. The white section for *P. sylvestris* in the Mediterranean biome represents missing data (due to its distribution in Spain).

### 3.2 Relative importance of climatic drought and basal area across latitude

In the case of Scots pine, basal area variables (i.e. *BAintra* and *BAinter*) were more important than drought-related variables (i.e. *WAI* and *SPEI*) in explaining the probability of mortality across the latitudinal gradient (Fig. 2a and Table 1). The conspecific basal area was the most important driver from south to north with a mean relative importance of 0.96 (Table 1). The order of importance of the four predictors was stable across latitude, except from 43° to 45° latitude (corresponding to the French part of the Mediterranean biome) where drought-related variables (mainly *SPEI*) were nearly as important as basal area variables (Fig. 2a). From south to north, high levels of both conspecific and heterospecific basal area and increases in drought intensity (i.e. low *SPEI*) were correlated with higher probability of mortality (Fig. 2a). In contrast, low *WAI* was associated with high mortality probabilities in the Mediterranean biome and with low mortality probabilities in the boreal biome (see changes from negative to positive influence in Fig. 2a).

**Table 1.** Mean relative importance of each predictor and mean annual predicted probability of both species mortality per biome. The relative importance of each variable was first computed for each tree from the logistic regression model (see section 2.5), by giving a value of one to the most influencing variable and scaling the remaining variables accordingly. Secondly, the relative importance and the annual predicted probabilities of mortality (*Pi* in the equation 1) were average for each biome. Numbers in brackets refer to 95% confidence intervals. *BAintra*: conspecific basal area (m^2^ ha^-1^), *BAinter*: heterospecific basal area (m^2^ ha^-1^), *WAI*: water availability index (adimensional), *SPEI*: Standardised Precipitation-Evapotranspiration Index (adimensional).

|  |  | *P. sylvestris* |  |  |  | *F. sylvatica* |  |  |
| --- | --- | --- | --- | --- | --- | --- | --- | --- |
|  |  | All biomes | Mediterranean biome | Cool temperate biome | Boreal biome | All biomes | Mediterranean biome | Cool temperate biome |
| Relative importance | *BAintra* | 0.96<br>(0.9616;<br>0.9630) | 0.95<br>(0.9455;<br>0.9482) | 0.96<br>(0.9625;<br>0.9644) | 0.99<br>(0.9860;<br>0.9870) | 0.61<br>(0.6134;<br>0.6162) | 0.64<br>(0.6411;<br>0.6465) | 0.61<br>(0.6076;<br>0.6107) |
|  | *BAinter* | 0.67<br>(0.6755;<br>0.6737) | 0.52<br>(0.5235;<br>0.5258) | 0.73<br>(0.7308;<br>0.7331) | 0.83<br>(0.8293;<br>0.8314) | 0.54<br>(0.5377;<br>0.5426) | 0.80<br>(0.7915;<br>0.8010) | 0.49<br>(0.4878;<br>0.4928) |
|  | *WAI* | 0.31<br>(0.3089;<br>0.3114) | 0.35<br>(0.3509;<br>0.3552) | 0.32<br>(0.3191;<br>0.3233) | 0.22<br>(0.2169;<br>0.2201) | 0.74<br>(0.7390;<br>0.7442) | 0.71<br>(0.6997;<br>0.7130) | 0.75<br>(0.7456;<br>0.7513) |
|  | *SPEI* | 0.44<br>(0.4370;<br>0.4397) | 0.40<br>(0.3976;<br>0.4017) | 0.42<br>(0.4218;<br>0.4258) | 0.53<br>(0.5265;<br>0.5311) | 0.70<br>(0.7018;<br>0.7066) | 0.61<br>(0.6055;<br>0.6168) | 0.72<br>(0.7196;<br>0.7249) |
| Annual predicted mortality |  | 0.0061<br>(0.00611;<br>0.00618) | 0.0077<br>(0.00763;<br>0.00775) | 0.0063<br>(0.00619;<br>0.00631) | 0.0033<br>(0.00332;<br>0.00337) | 0.0038<br>(0.00374;<br>0.00382) | 0.0052<br>(0.00506;<br>0.00530) | 0.0035<br>(0.00347;<br>0.00354) |

**Figure 2.**
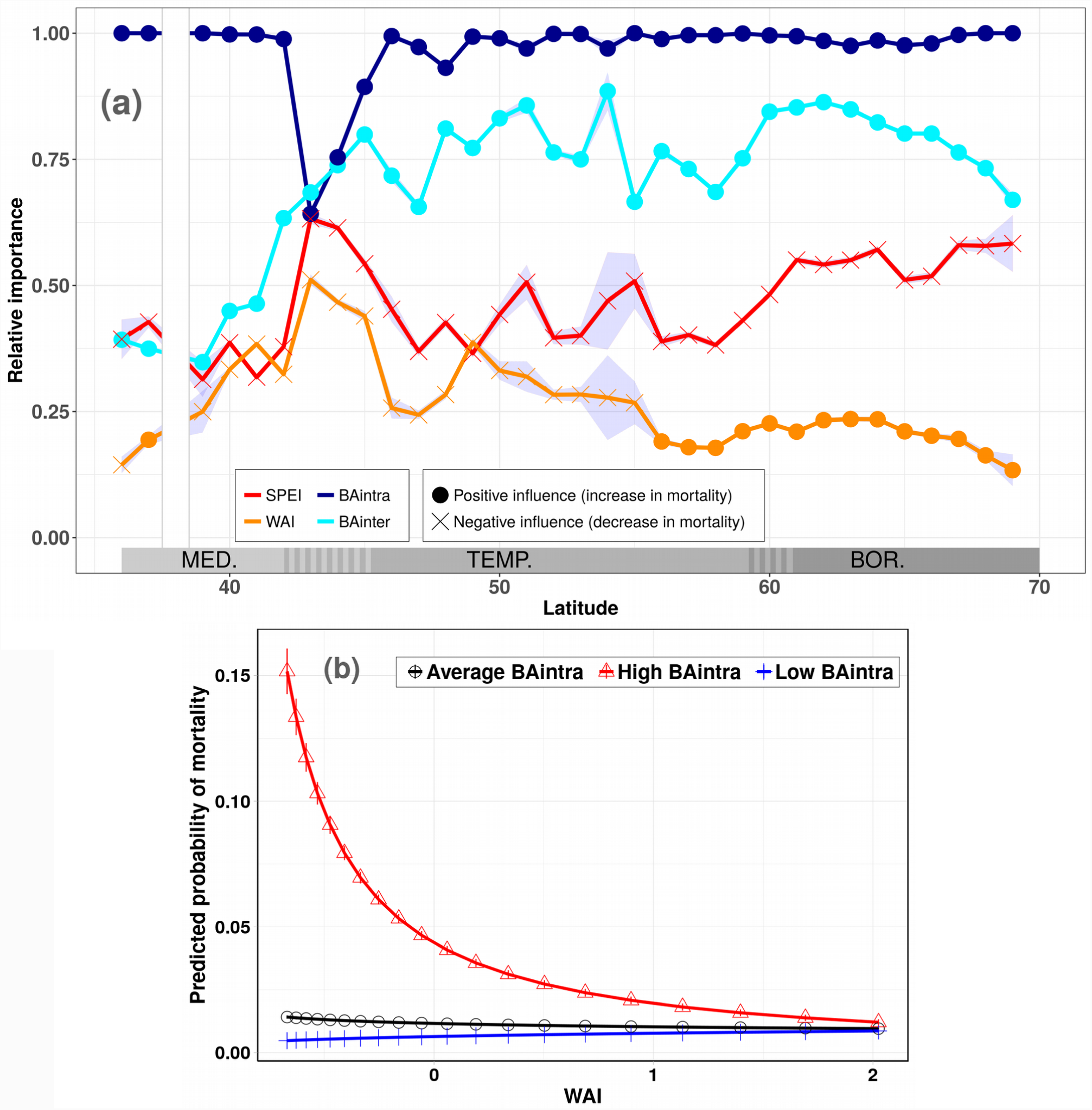
Effects of drought-related variables and basal area on Scots pine mortality. a) Relative importance of the changes in climatic drought intensity over the study period (i.e. *SPEI*), climatic drought intensity (i.e. *WAI*), conspecific basal area (i.e. *BAintra*) and heterospecific basal area (i.e. *BAinter*) on Scots pine predicted probability of mortality. The relative importance of each variable was computed for each tree from the logistic regression model (see section 2.5), by giving a value of one to the most influencing variable and scaling the remaining variables accordingly. For each variable, the relative importance values were aggregated by 1° latitude resolution and the points of the graph correspond to the average values. The grey areas around the curves correspond to the 95% confidence intervals. The acronyms MED., TEMP. and BOR. in grey bars refer to the Mediterranean, cool temperate and boreal biome, respectively. The white section corresponds to missing data at that latitude (due to Scots pine distribution in Spain). **b) Interactions between conspecific basal area (i.e. *BAintra*) and climatic drought intensity (i.e. *WAI*) on Scots pine probability of mortality.** This interaction was considered significant if its z value was lower than -2 or higher than 2 and was the most important interaction influencing Scots pine mortality (Table S3.1). Scots pine mortality was predicted at three different levels of conspecific basal area (mean value, 99.5 ^th^ percentile and 0.005^th^ percentile; proxies of average, high and low competition, respectively) along a drought gradient while the other predictors were fixed at their mean value.

For beech trees, drought-related variables were more important than basal area variables in explaining mortality probability across the major part of the latitudinal gradient (except in the south) with a mean relative importance of 0.74 and 0.70 for *SPEI* and *WAI*, respectively (Fig. 3a and Table 1). Low *WAI* and *SPEI* were associated with higher mortality rates (see negative influence in Fig. 3a). The relative importance of conspecific basal area remained stable across latitude whereas that of heterospecifics varied from being the most important variable explaining beech mortality in the Mediterranean biome to being the least important one in the cool temperate biome (Fig. 3a and Table 1). Beech mortality probability increased with conspecific basal area and decreased with heterospecific basal area (Fig. 3A and Table 1).

**Figure 3.**
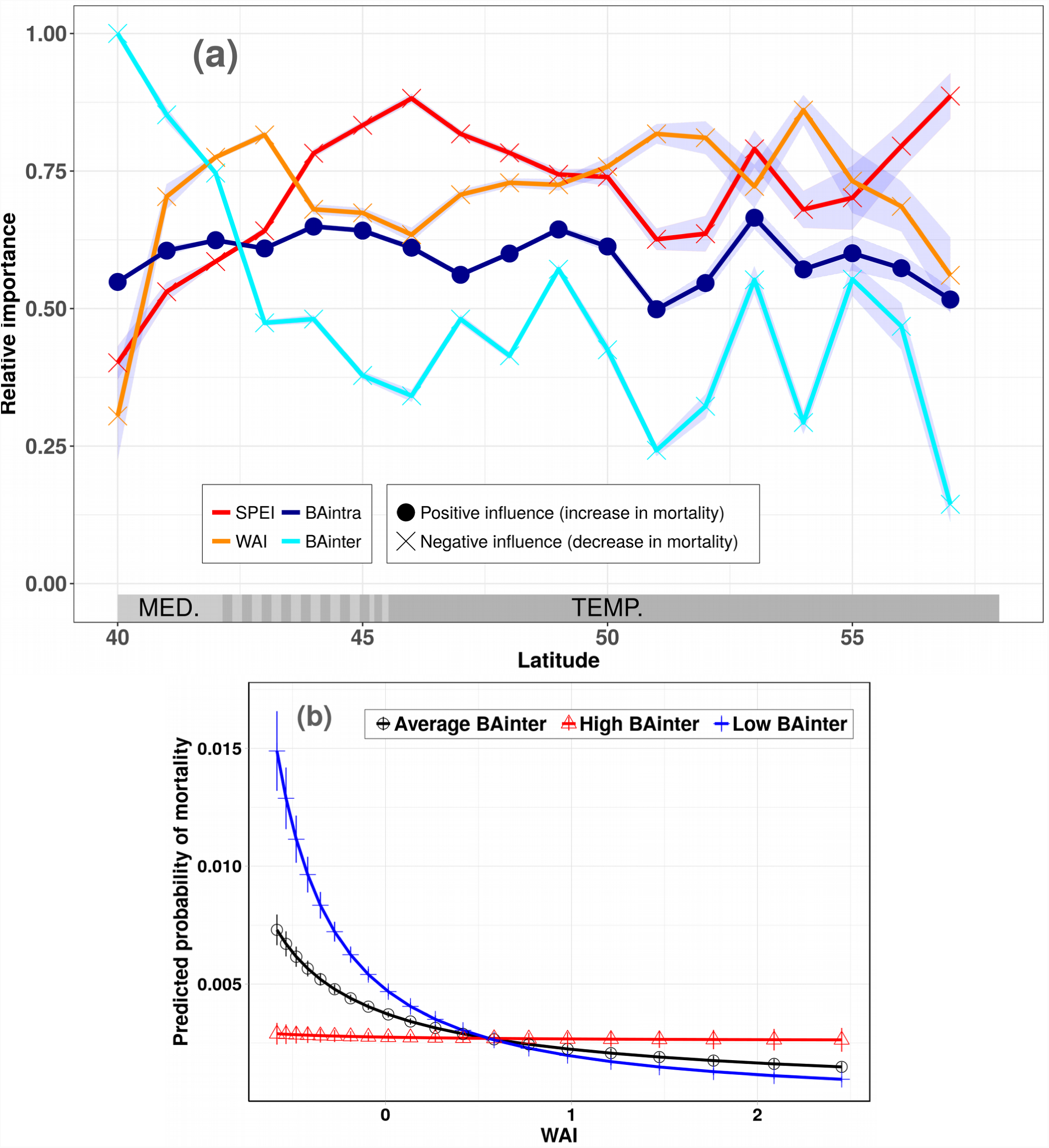
Effects of drought-related variables and basal area on beech mortality. a) Relative importance of the changes in climatic drought intensity over the study period (i.e. *SPEI*), climatic drought intensity (i.e. *WAI*), conspecific basal area (i.e. *BAintra*) and heterospecific basal area (i.e. *BAinter*) on beech predicted probability of mortality. The relative importance of each variable was computed for each tree from the logistic regression model (see section 2.5), by giving a value of one to the most influencing variable and scaling the remaining variables accordingly. For each variable, the relative importance values were aggregated by 1° latitude resolution and the points of the graph correspond to the average values. The grey areas around the curve correspond to the 95% confidence intervals. The acronyms MED. and TEMP. in grey bars refer to the Mediterranean and cool temperate biome, respectively. **b) Interaction between heterospecific basal area (i.e. *BAinter*) and climatic drought intensity (i.e. *WAI*) on beech probability of mortality.** This interaction was considered significant as its z value was higher than 2 (see Table S3.1). Beech mortality was predicted at three different levels of heterospecific basal area (mean value, 99.5^th^ percentile and 0.005^th^ percentile; proxies of average, high and low competition, respectively) along a drought gradient while the other predictors were fixed at their mean value.

### 3.3 Interactions between climatic drought and basal area

In the Scots pine model, all interactions between drought-related variables (i.e. *WAI* and *SPEI*) and basal area variables (i.e. *BAintra* and *BAinter*) were significant (Table S3.1). The strongest interaction was between climatic drought intensity and conspecific basal area (i.e. *WAI* and *BAintra*; Table S3.1): regardless of drought intensity, the probability of mortality remained weak when the conspecific basal area was low or intermediate, whereas it strongly increased in dry areas where the conspecific basal area was high (Fig. 2b; see Fig. S4 for the other interactions that affected mortality weakly, albeit significantly).

In the beech model, the only significant interaction was that between climatic drought and heterospecific basal area (*WAI* and *BAinter*; Table S3.1): the probability of mortality increased in dry areas where heterospecific basal area was low or intermediate, while the probability of mortality remained stable (and always low) in dry areas where heterospecific basal area was high (Fig. 3b).

### 3.4 Spatial patterns of predicted tree mortality across Europe

Across their range, the predicted annual probability of Scots pine mortality was on average higher than that of beech (0.0061 and 0.0038, respectively; Table 1) but followed the same trend across the latitudinal gradient (Fig. 4). The highest predicted mortality rates for both species were in south-eastern France, at the ecotone between the Mediterranean and cool temperate biomes (Fig. 4).

**Figure 4.**
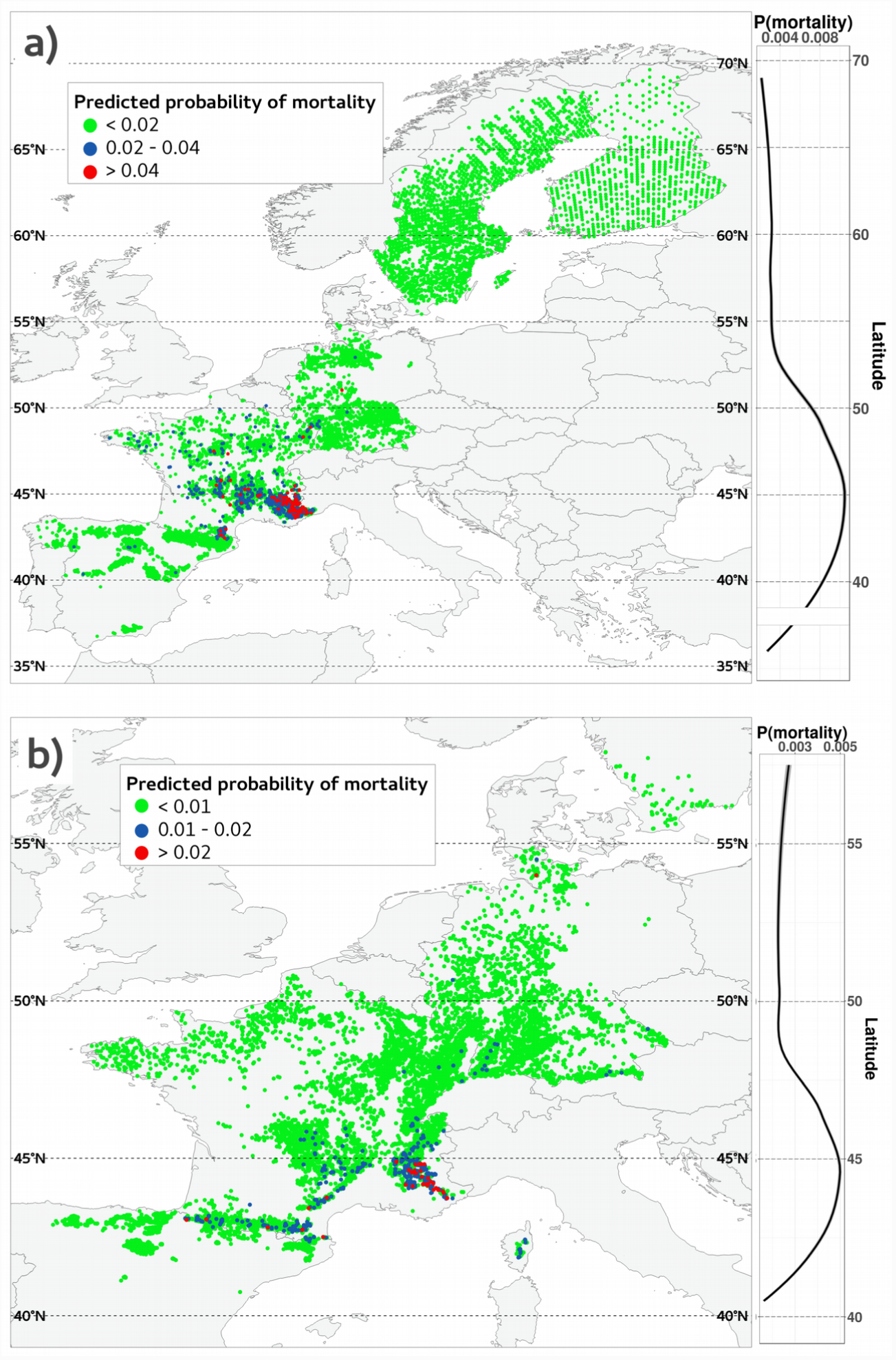
Spatial projection of the annual predicted probability of mortality at the individual-level across Europe for a) *P. sylvestris* and b) *F. sylvatica*. Graphs in the right panels display predictions (noted as *P(mortality)*) across latitude. For both species, predictions were calculated for all trees from the logistic regression model and were clustered at 1° latitude for *P. sylvestris* and at 0.5° resolution for *F. sylvatica*. A locally weighted regression was used to obtain the smooth solid lines (“loess” method of the geom_smooth function in “ggplot2” R package). Grey areas indicate 95% confidence intervals (almost confused with the curves). The white section for *P. sylvestris* in the Mediterranean biome represents missing data at that latitude (due to its distribution in Spain).

The predicted rates of Scots pine mortality were highest in the Mediterranean biome (mean value of 0.0077 for 62,165 trees), intermediate in the cool temperate biome (mean value of 0.0063 for 62,914 trees) and lowest in the boreal biome (mean value of 0.0033 for 36,641 trees) (Table 1). Similarly, the predicted individual probability of beech mortality was higher in the Mediterranean biome (mean value of 0.0052 for 9,315 trees) than in the cool temperate biome (mean 0.0035 for 47,876 trees) (Table 1).

## 4 DISCUSSION

Exploring the drivers of background tree mortality at a continental scale opens a new perspective for understanding tree mortality patterns across species’ ranges, including some demographic events observed at a smaller scale (Carnicer *et al.*, 2011). Although considerable attention has been paid to the effects of drought and basal area on tree mortality (Mantgem *et al.*, 2009; Greenwood *et al.*, 2017; Hember *et al.*, 2017; Ruiz-Benito *et al.*, 2013), our results demonstrate that the combination of the two, through direct and indirect effects that vary along geographical gradients and between the two species (Fig. 2 and 3), is shaping background mortality across species’ ranges (see also Ruiz-Benito et al., 2013; Jump et al., 2017; Young et al., 2017). Interestingly, both species had similar patterns of predicted mortality, with the highest mortality rates in the southern French part of the Mediterranean biome (Fig. 4).

### 4.1 Increase in climatic drought intensity associated with higher mortality rates

Drought-related variables were more important for beech mortality than Scots pine (Fig. 2 and 3), probably reflecting functional differences in species responses to drought (Choat *et al.*, 2018). Scots pine is a drought-avoiding species that can survive from wet to dry environments, whereas beech is a drought-sensitive species. Nevertheless, both beech (i.e. an angiosperm and broad-leaved species) and Scots pine (i.e. a gymnosperm and evergreen species) exhibited higher mortality rates in areas that were subject to increasing droughts during the study period (negative *SPEI*; Fig. 2 and 3). This result suggests that major phylogenetic and functional groups could display a similar mortality response to increasing drought (Greenwood *et al.*, 2017) and is consistent with the results of a multi-species study suggesting that climatic extremes (like extreme droughts) are affecting tree mortality in Europe (Neumann *et al.*, 2017).

The increase in drought intensity that occurred at about 45° latitude during the study period (see the lowest *SPEI* values in Fig. S2.1 and S2.3b) could be responsible for the higher tree mortality rates in the Mediterranean biome (Fig. 4), which is also supported by the high relative importance of the increase in drought intensity at this latitude (see the highest values of *SPEI* in Fig. 2 and 3). Moreover, we observed higher mortality rates in the driest areas (i.e. low *WAI*), as already reported for Scots pine in some inner Alpine valleys (Rigling *et al.*, 2013) and in the Iberian Peninsula (Vilà-Cabrera *et al.*, 2011; Galiano *et al*., 2011). Nevertheless, the stronger effect of increasing droughts over the study period (i.e. *SPEI*) than that of drought intensity (i.e. *WAI*) on Scots pine mortality could mean that mortality events tend to occur when drought conditions exceed the average in a given area, suggesting a certain degree of Scots pine adaptation to local conditions (Savolainen *et al.*, 2007).

Drought-related variables were key drivers of beech mortality and were comparatively more important than heterospecific and conspecific basal area. A regional study of tree mortality suggested that competition between trees is more important than climate (Ruiz-Benito *et al.*, 2013), but that study did not cover a climatic gradient as large as our study. Continental-scale studies of beech tree-level mortality in response to drought are almost no-existent, and growth-based studies produced contradictory results showing both drought-induced reduction in growth (Jump *et al.*, 2006) and drought-associated increase in growth (Tegel *et al.*, 2014). However, beech mortality responses to changes in drought intensity may differ from growth responses as this species can survive low growth periods before death (Hülsmann *et al.*, 2018).

### 4.2 Conspecific and heterospecific neighbours can affect individual tree mortality differently

Competition is a critical driver of forest structure (Kunstler *et al.*, 2016), which strongly influences tree mortality and is comparatively more important for shade-intolerant than shade-tolerant species (Ruiz-Benito *et al.*, 2013). High mortality rates were associated with high conspecific basal area in both species and high heterospecific basal area in Scots pine. However, high heterospecific basal area was correlated with low mortality rates in beech (Fig. 2 and 3). Scots pine is a shade-intolerant tree which is highly sensitive to competition for light (Ruiz-Benito *et al.*, 2013), which might explain why both intra and inter-specific competition strongly and positively influenced its mortality rate (Condés & del Río, 2015). In contrast, beech is a late successional and shade-tolerant species (Hülsmann *et al.*, 2018) that outcompetes other species in fertile sites (Condés & del Río, 2015). This is consistent with our observation of high mortality rates with high conspecific basal area but also with low heterospecific basal area: beech mainly suffers from the presence of conspecific neighbours, but not from heterospecific neighbours, which are necessarily less competitive species. This result is supported by growth studies showing that beech benefits from admixture with other species but is highly sensitive to intra-specific competition (Ratcliffe *et al.*, 2015).

The heterospecific basal area affected the mortality rates of both species less than the conspecific basal area (Table S3.1, Fig. 2 and 3). The dominant nature of both Scots pine and beech in European forests may partly explain this difference as the basal area of heterospecifics was much lower than that of conspecifics all along the latitudinal gradient (Fig. S2.1). Nevertheless, the overdominance of intra-specific competition, a key process for stabilising ecosystems, is a globally-observed pattern (Kunstler *et al.*, 2016), which could be linked to how interspecific differences determine complementarity mechanisms and, consequently, individual resource-use and coexistence mechanisms (Ruiz-Benito et al., 2017b).

### 4.3 The effects of climatic drought and basal area should be considered jointly in mortality studies

Competition with neighbours can be expressed as asymmetric competition for light on small suppressed trees (Ruiz-Benito *et al.*, 2013) but also as symmetric competition for limited resources, like water or nutrients (Franklin *et al.*, 1987). Drought-induced mortality may be strong in areas with high levels of competition, because plants are more stressed and small changes in water availability could result in massive mortality events (Ruiz-Benito *et al.*, 2013; Young *et al.*, 2017). In the case of Scots pine, the strong interaction between drought intensity and conspecific basal area reinforces this assumption (Table S3.1). Indeed, mortality rates were high in dry areas with high conspecific basal area whereas in areas with lower conspecific basal area, trees had still sufficient resources to survive despite reduced water availability (Fig. 2b). This result suggests that Scots pine suffers from the presence of neighbouring trees only when resources are scarce (Young *et al.*, 2017).

In the case of beech, the influence of conspecific basal area on mortality was not modulated by drought (Table S3.1), suggesting that resource depletion does not exacerbate competitive pressure among beech trees. However, in the driest areas, the probability of beech mortality was considerably higher when heterospecific basal area was low (Fig. 3b), suggesting that beech benefits from the presence of other species. In contrast, in the wettest sites, the probability of beech mortality was constant regardless of basal area values (Fig. 3b). This pattern may result from the stress-gradient hypothesis: facilitation prevails in water-limited areas while competition in non-limiting conditions (Callaway & Walker, 1997). Beech may be highly favoured by facilitation because of its shade-tolerance and recent works have highlighted that beech trees are more resilient and resistant to drought in mixed stands (i.e. particularly with oaks; Pretzsch et al., 2012). One possible explanation is that beech benefits from the hydraulic lift of water by oaks (i.e. release of water from the deep root-system of oaks in the upper layer of soils; Zapater et al., 2011). This may explain why heterospecific basal area was the most important predictor in the Mediterranean biome (Fig. 3 & Table 2): beech needs its neighbours to survive in the Mediterranean region.

Previous regional studies reported contradictory interaction effects between competition and drought, for instance for Scots pine: higher rate of decline in dry areas but only at low competition levels (Vilà-Cabrera *et al.*, 2013), low mortality rates related to high heterospecific basal area in wet areas (Condés & del Río, 2015) and only additive effects of competition and drought on mortality with no interaction effects (Galiano *et al.*, 2010). Our study is the first to describe interaction patterns between drought and basal area at the scale of the distribution of each species (Fig. 2, 3 and S4). As we found four significant interactions (albeit three of which only slightly affect mortality) influencing Scots pine mortality and only one in the case of beech (Table S3.1), we can assume that Scots pine mortality is affected directly and indirectly by drought through interactions with basal area while beech mortality was more directly affected by drought.

### 4.2 Tree mortality patterns along latitude and potential associated range shifts

Mortality rates of both beech and Scots pine are higher in the southern part of their distribution, mainly corresponding to the French part of the Mediterranean biome and the Pyrenees in the case of beech (Fig. 4). As the Scots pine model underestimated the probability of mortality in the extreme parts of the latitudinal gradient (Fig. 1a), the decrease in mortality might be less sharp in the Mediterranean biome and there may not be a slight decrease at the northern end. On the contrary, beech model overestimated the probability of mortality in the southern part of the gradient (Fig. 1b) and the northern half of the cool temperate biome (Fig. 4), suggesting that the peak of the probability of mortality in the southern half of the cool temperate biome is sharper than predicted.

An unexpected result was that French Mediterranean Scots pines and beech trees suffered even more from climatic drought than those in Spain, where several studies reported high mortality or defoliation rates in the Iberian Peninsula in both species (Carnicer *et al.*, 2011; Vilà-Cabrera *et al.*, 2011, 2013; Benito Garzón *et al.*, 2013). Nevertheless, this pattern may be explained by the high altitudes at which both species occur in Spain, and the calcareous soils of southeastern France, which do not retain water and are consequently very dry. In the case of Scots pine, we also hypothesise that local adaptation to temperature explains our underestimated mortality predictions in the southernmost part of the gradient (Savolainen *et al.*, 2007): populations in these areas may be highly locally-adapted to drought conditions and therefore less resistant to changing climate (Benito Garzón *et al.*, 2011).

The high mortality rates predicted in the French part of the Mediterranean biome could be explained by the increase in drought intensity during the study period in that region (Fig. S2.3b), suggesting that mortality plays a critical role in delimiting the driest part of the species ranges (Gaston, 2009; Benito Garzón et al., 2013; Ruiz-Benito et al., 2017a), in particular in the Mediterranean biome, which is expected to face drier conditions in the coming decades. In addition to direct effects of climate change, Scots pine and beech are exposed to more intense fires in the driest parts of their range (Fréjaville *et al.*, 2018) and these should increase the likelihood of range contraction at the ecotone between Mediterranean and cool temperate biomes.

### 4.3 Limitations

Until recently, European forests have been extensively exploited and forest management is still widespread, particularly in the Scandinavian countries (Schelhaas *et al*., 2018). Although we removed the direct effects of management in our study (i.e. by removing plots in which trees were noted as harvested), management may still result in both an overestimation (e.g. by reducing competition pressure in thinned plots) and an underestimation of natural mortality rates through salvage loggings (i.e. the harvest of dead trees after a natural disaster) or sanitation fellings (i.e. the harvest of diseased trees).

Other factors also affect tree mortality: insect outbreaks (Anderegg *et al.*, 2015), mistletoe (Dobbertin & Rigling, 2006), atmospheric pollutants (Dietze & Moorcroft, 2011), populations genetic differentiation and plasticity (Benito Garzón *et al.*, 2011), soil characteristics (Dietze & Moorcroft, 2011). However, given our concern to limit the model complexity and the lack of large-scale data, we decided not to include them in our study and to focus on comparing the effects of drought and competition on mortality.

## CONCLUSIONS

Mortality of Scots pine and beech was affected by drought and competition, but in different ways. Drought directly affected beech mortality rates and beech trees benefited from mixing with other species, particularly in the Mediterranean biome. Scots pine mortality suffered mostly from competition and was indirectly affected by drought through interactions with competitors, especially in southeastern France. In this area, which experienced a marked increase in drought intensity during the study period, high mortality rates were predicted for both species, as expected for temperate trees for which the Mediterranean biome corresponds to the southernmost part of the distribution. In a warming climate, our study is a step further in understanding geographical patterns of tree mortality in Europe and shed light on the high mortality risks faced by European tree species, regardless of their different life-history strategies, especially at the ecotone between the Mediterranean and cool temperate biomes. In this priority area, beech could benefit from mixing with other species and pine from reduced competition.

## DATA ACCESSIBILITY

The data are available upon request to the co-authors.

## ACKNOWLEDGEMENTS

This study was funded by the “Investments for the Future” program IdEx Bordeaux (ANR-10-IDEX-03-02) and the FunDivEUROPE project was supported by the European Union Seventh Framework Programme (FP7/2007-2013) under grant agreement no 265171. MAZ and PRB were also supported by project FUNDIVER (Spanish Ministry of Economy and Competitiveness, MINECO, Spain; CGL2015-69186-C2-2-R). We thank Christian Wirth and Gerald Kaendler for access to the FunDivEUROPE data, the MAGRAMA for access to the Spanish Forest Inventory, the Johann Heinrich von Thünen-Institut for access to the German National Forest Inventories, the Natural Resources Institute Finland (LUKE) for making permanent sample plot data from the Finnish NFI available, the Swedish University of Agricultural Sciences for making the Swedish NFI data available. We thank Georges Kunstler for his help in developing the models and the Forest Ecology and Restoration group of the University of Alcalá. We also thank Frédéric Barraquand for the discussions about the model equations.

