## Supplementary Information for "Similar patterns of background mortality across Europe are mostly driven by drought in European beech and a combination of drought and competition in Scots pine"

### SUPPORTING INFORMATION

**Appendix S1.** Further details on national forest inventories.

**Appendix S2.** Fluctuations of basal area and drought-related variables across latitude.

**Appendix S3.** Further details on model selection and validation.

**Appendix S4.** Interactions between basal area and drought-related variables not shown in the main figures for Scots pine.

### Appendix S1. Further details on national forest inventories (NFIs).

To harmonise the data between NFIs, we removed all trees with a diameter at breast height (*DBH*, cm) less than 10 cm and all plots with a total basal area less than 4 $m^2$ $ha^{-1}$. We also removed plots where trees had been harvested during the period between the surveys, to avoid any bias due to forest management. However, we note that there could still be an influence from other management activities.

#### NFIs protocols of Germany, Finland, Spain and Sweden

NFIs were harmonised within the FunDivEUROPE project (Baeten *et al.*, 2013) and were on permanent plots sampled from 1981 to 2005 for the first survey and from 1990 to 2011 for the second one. For details on each NFI protocol, see Appendix S1 of Ratcliffe *et al.* (2016).

Regarding the tree mortality census in these countries, a tree was considered dead if it was alive in the first survey but was found dead in the second survey.

#### French NFI protocol

We used data from the temporary plots of the 2005-2014 annual campaigns of French NFI. Sample plots are on a systematic 1 $km^2$ grid in forested areas of the country (IGN Inventaire Forestier; http://inventaire-forestier.ign.fr/spip/spip.php?rubrique153). The French NFI used a variable radius plot size depending on the *DBH* (diameter at breast height) of the sample trees; each plot has three nested subplots of 6, 9 and 15 m radius and the minimum *DBH* for a tree to be recorded within a subplot is 7.5 cm, 22.5 cm and 37.5 cm, respectively. A relative area weight is attributed to each inventoried tree to obtain a per ha quantities. The readers are referred to the study of Charru *et al.* (2010) for a more detailed description of French NFIs. A tree was noted as dead if its date of death was estimated by direct observation at less than five years; blow down trees were not considered in the present study.

**Figure S1.1. National forest inventory plots sorted by biome for (a) *P. sylvestris* and (b) *F. sylvatica*.** The classification follows Olson *et al.* (2001).

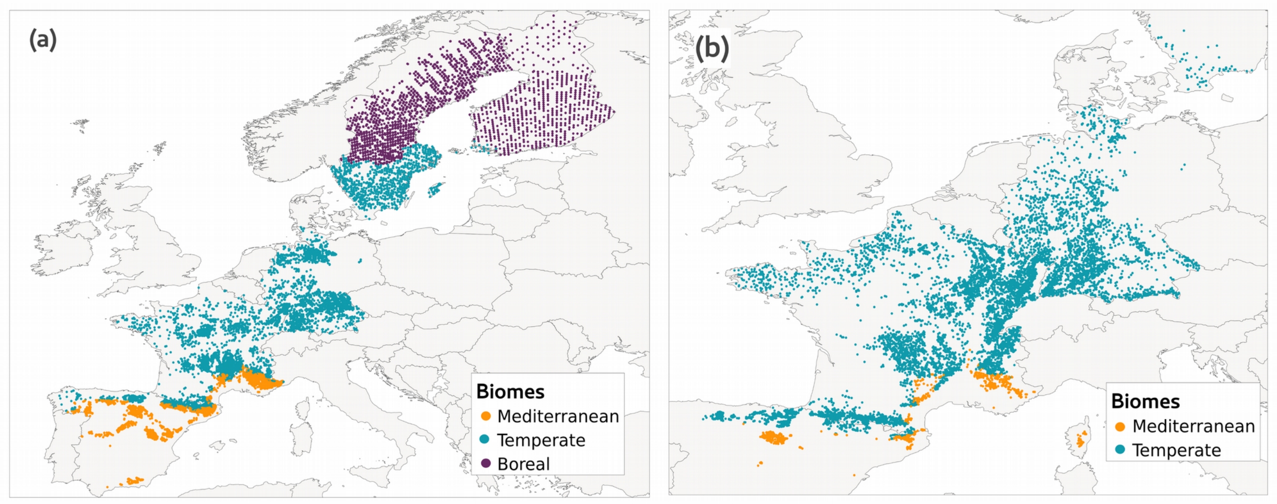


**Figure S1.2. Number of national forest inventory plots along the latitudinal gradient for (a) *P. sylvestris* and (b) *F. sylvatica*.**

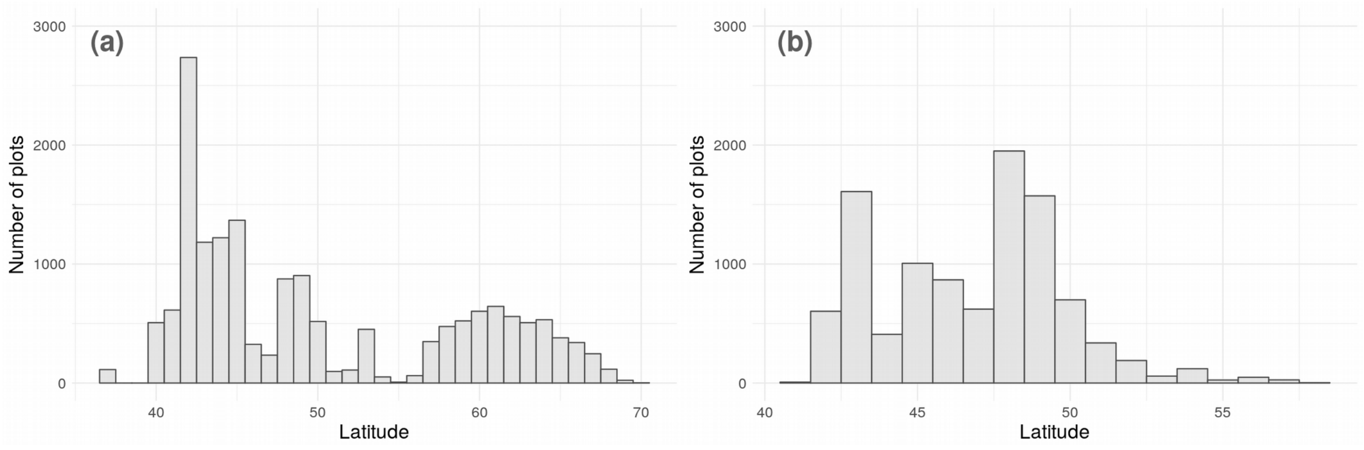

#### Appendix S2. Fluctuations of basal area and drought-related variables across latitude.

Drought-related variables (i.e. *WAI* and *SPEI*) showed similar trends across the latitudinal gradient: high values in the boreal and the major part of the cool temperate biome (i.e. corresponding to areas that did not experience climatic droughts), a sharp decrease in the French part of the Mediterranean biome and the lowest values in the Mediterranean biome, indicating severe drought (Fig. S2.1 and Table S2.1).

In the case of Scots pine, plot-level basal area of all species (i.e. *BAall*) had the lowest values at the northernmost part of the range (down to 10 $m^2$ $ha^{-1}$) and the highest values from the southern half of the cool temperate biome to the northern part of the Mediterranean biome (up to 30 $m^2$ $ha^{-1}$, Fig. S2.2a). For beech, plot-level basal area of all species varied between 20 $m^2$ $ha^{-1}$ and 35 $m^2$ $ha^{-1}$, with the highest values at about 45° latitude and to a lesser extent, at about 52° latitude (Fig. S2.2b). For both species, conspecific basal area (i.e. *BAintra*) was higher than heterospecific basal area (i.e. *BAinter*) along the latitudinal gradient. This was especially strong in Scots pine in the Mediterranean biome where plots were almost monospecific (i.e. *BAintra* nearly equal to *BAall*, Fig. S2.2a).

**Figure S2.1. Raw values of *SPEI* and *WAI* along the European latitudinal gradient covered by national forest inventory plots within which at least one individual of Scots pine or European beech was sampled.** Climatic values were aggregated by 1° latitude resolution and graph curves correspond to the average values. Red areas depict 95% confidence intervals (almost confused with the curve in the case of *WAI*). The acronyms MED., TEMP. and BOR. in grey bars refer to the Mediterranean, cool temperate and boreal biome, respectively. The white section corresponds to missing data in Scots pine dataset.

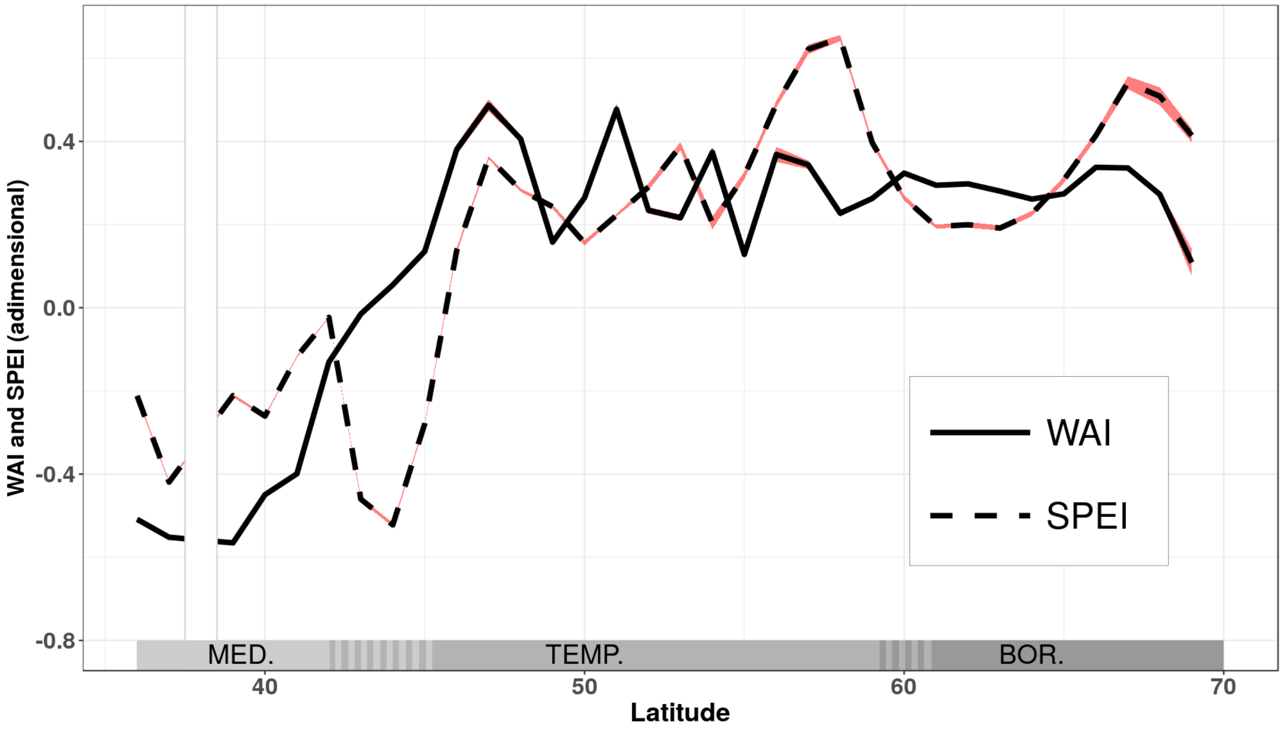


**Table S2.1. Average of *WAI* and *SPEI* raw values within each biome during the study period in national forest inventory plots including at least one individual of our two species of interest.** Values in brackets correspond to the lower and upper endpoint of the 95% confidence interval, respectively.

| | | ***WAI*** (adimensional) | ***SPEI*** (adimensional) |
|---|---|---|---|
| **Biomes** | **Mediterranean** | -0.30<br>(-0.2995, -0.3022) | -0.21<br>(-0.2123, -0.2162) |
| | **Cool temperate** | 0.15<br>(0.1476, 0.1515) | 0.07<br>(0.0657, 0.0701) |
| | **Boreal** | 0.31<br>(0.3093, 0.3118) | 0.29<br>(0.2841, 0.2891) |

**Figure S2.2 Raw values of the total basal area (*BAall*, $m^2$ $ha^{-1}$), the conspecific basal area (*BAintra*, $m^2$ $ha^{-1}$) and the heterospecific basal area (*BAinter*, $m^2$ $ha^{-1}$) along the European latitudinal gradient in national forest inventory plots within which at least one individual of Scots pine (a) or European beech (b) was sampled.** Values were aggregated by 1° latitude resolution for both species and graph curves correspond to the average values. Grey areas depict 95% confidence intervals. The acronyms MED., TEMP. and BOR. in grey bars refer to the Mediterranean, cool temperate and boreal biome, respectively. The white section corresponds to missing data in Scots pine dataset.

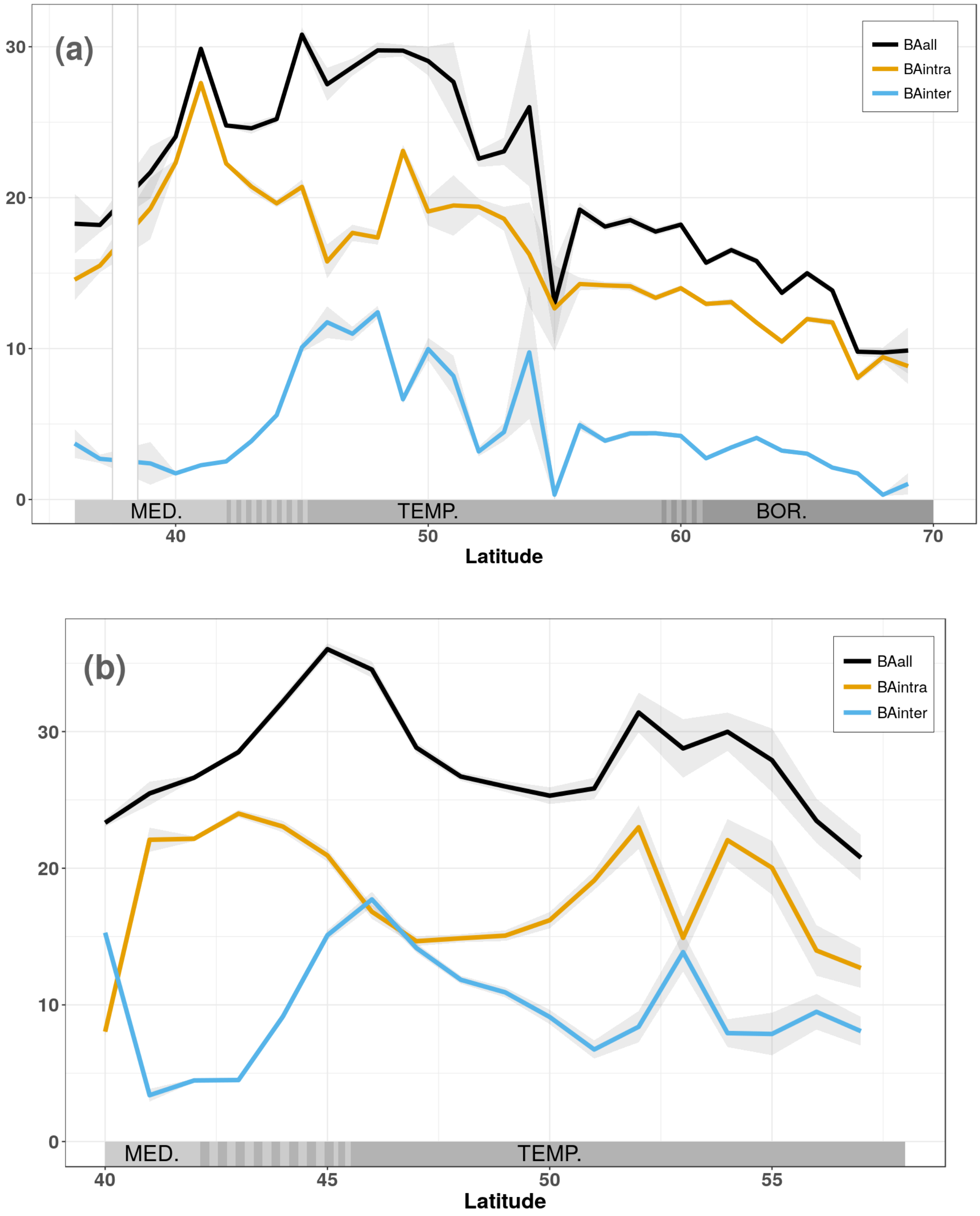

**Figure S2.3 Map of the predictor's raw values in each national forest inventory plot.** (a) *WAI* (adimensional) for both species. (b) *SPEI* (adimensional) for both species. (c) and (d) *BAintra* and *BAinter* ($m^2$ $ha^{-1}$) for Scots pine. (e) and (f) *BAintra* and *BAinter* ($m^2$ $ha^{-1}$) for common beech.

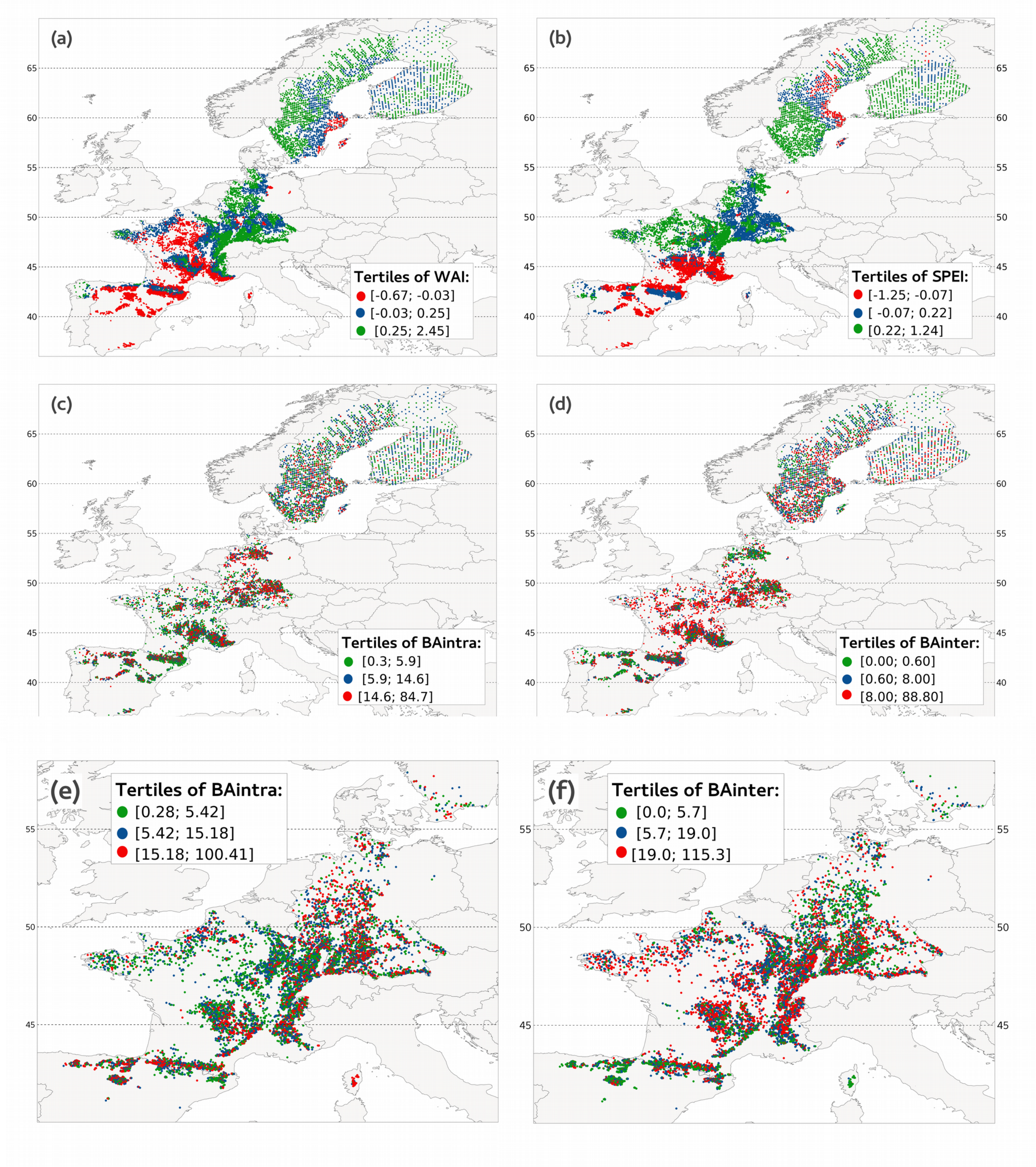

#### Appendix S3. Further details on model selection and validation.

##### Selection of the transformation of each variable

To select the best transformation for each variable, similar for both species, we followed these steps:

1- We created a common dataset that combined the whole beech’s dataset and a sample of Scots pine’s dataset with size corresponding to the beech dataset's size (so that the numerical superiority of Scots pines didn’t bias the choice of each variable transformation).

2- Each variable *x* was transformed in $x^2$ and in log($x+c$), with *c* a variable-specific value allowing to account for negative values, if needed.

3- Each variable *x* and its transformations log($x+c$) and $x^2$ were standardised (i.e. subtracted the mean value and divided by the standard deviation).

4- For each variable *x*, we ran the model described in the equation 1 on the common dataset and with the function *k* successively replaced by one of the three transformations: *x*, $x^2$ or log($x+c$). We kept the transformation corresponding to the lowest Bayesian Information Criterion (BIC; Schwarz, 1978) for the following analyses, namely: *BAintra*, log(*BAinter*), log(*WAI*) and log(*SPEI*). The partial residual plots confirmed that with the transformation applied we meet the linearity of the *xy* relationship (Figure S3.2).

##### *DBH’s* calculation

*DBH* (diameter at breast height) can influence tree mortality through a U-shape (Lines *et al.*, 2010) or a L-shape (Ruiz-Benito *et al.*, 2013) relationship with mortality. For that reason, the influence of tree size on tree mortality is often modelled as $a_{sp} \times DBH + b_{sp} \times \log(DBH)$. As we required a single parameter to estimate the relative importance of each predictor (see section 2.5), we calculated a non-linear variable from *DBH*:

$$DBHnl_{sp,i} = DBH_i + r_{sp} \times \log(DBH_i) \qquad (4)$$

which is equivalent to $a_{sp} \times DBH + b_{sp} \times \log(DBH)$, where $r_{sp} = b_{sp}/a_{sp}$ is a species-specific parameter and $DBH_i$ is the *DBH* of each individual tree *i*.

For each species separately, we estimated $r_{sp}$ by replacing k in equation 1 with $a_{sp} \times DBH + b_{sp} \times \log(DBH)$, and then fixing $r_{sp}$ in the final model run. The $r_{sp}$ values obtained and fixed in the final model were (with the 95% confidence interval in brackets):

$r_{pinus}$ = - 2.07 (-2.26; -1.88) and $r_{fagus}$ = -1.43 (-1.59; -1.27).

#### Sensitivity analysis

We conducted a sensitivity analysis to ensure that the estimated value of $r_{sp}$ (see equation 4 section "DBH calculation" above) that we used to calculate $DBHt_{sp}$ did not influence our results. To that end, we repeated the analysis where $r_{sp}$ was replaced by the lower and the upper endpoint of its 95% confidence interval. We used $SEa_{sp}$ and $SEb_{sp}$ the standard errors of $a_{sp}$ and $b_{sp}$ to calculate $SEr_{sp}$ the standard error of $r_{sp}$:

$$SEr_{sp} = (1 / b_{sp}) \times SEa_{sp} - (a_{sp} / b_{sp}^2) \times SEb_{sp}$$

The lower and upper endpoints of the 95% confidence interval of $r_{sp}$ were computed for both species as follows:

$$r_{lwr/upr,sp} \sim r_{sp} +/- SEr_{sp} \times 1.96$$

Then, we replaced $r_{upr,sp}$ and $r_{lwr,sp}$ in the final model for each species, recalculated each variable relative importance and plotted it against latitude. The patterns are robust along the entire latitudinal gradient (Fig. S3.1 and S3.2).

**Figure S3.1 Relative importance of drought-related variables and basal area for *F. sylvatica* probability of mortality.** The coefficient $r_{sp}$ was replaced by the upper (a) and the lower (b) endpoint of its 95% confidence interval. The relative importance was computed for each tree from the logistic regression model. The relative importance of the most influencing variable had a value of one and that of the other variables was scaled accordingly. For each variable, the relative importance values were aggregated by 1° latitude resolution and the points of the graph correspond to the average values. The grey areas around the curve correspond to the 95% confidence intervals. The acronyms MED. and TEMP. in grey bars refer to the Mediterranean and the temperate biome, respectively.

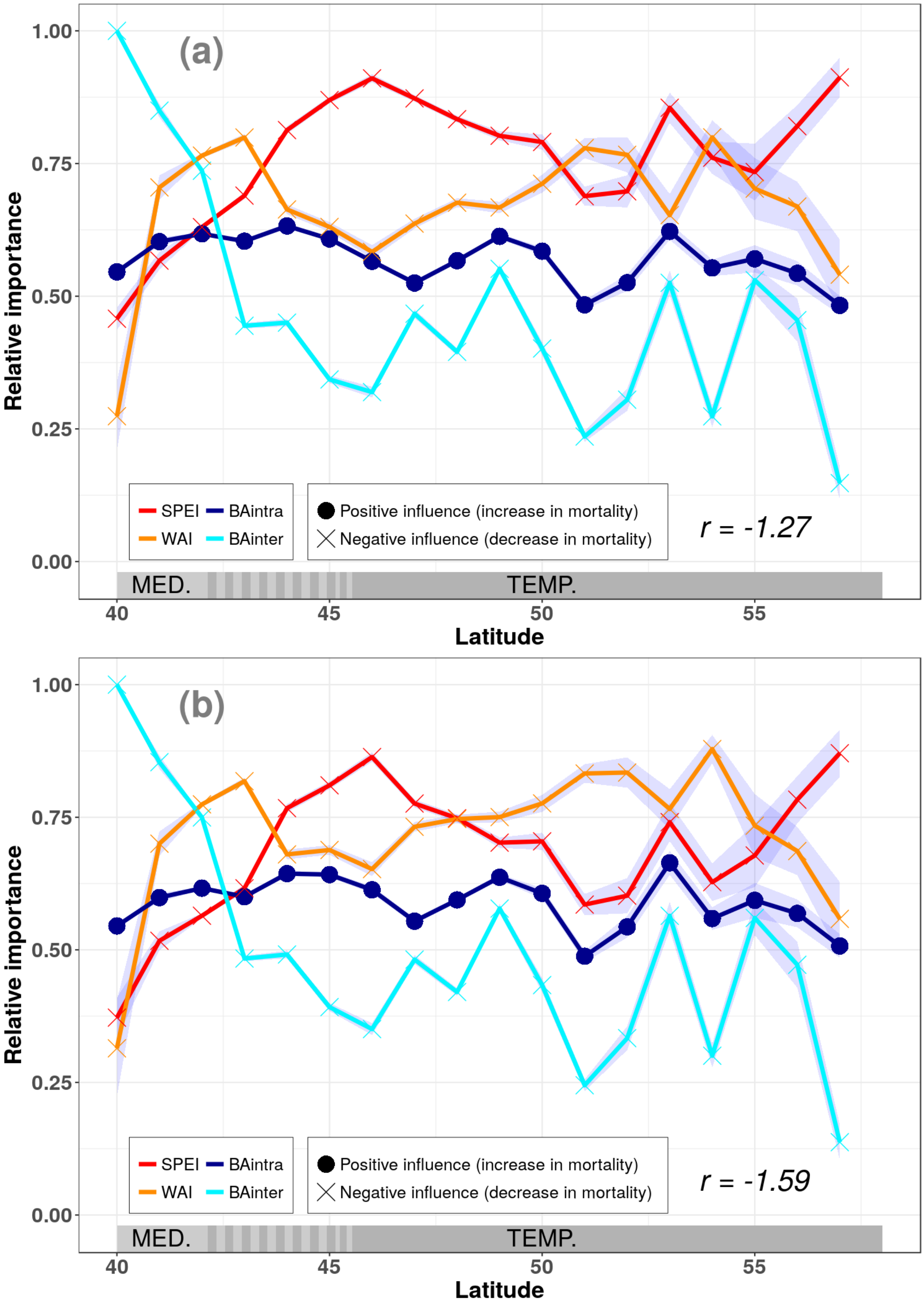

**Figure S3.2 Relative importance of drought-related variables and basal area for *P. sylvestris* probability of mortality.** The coefficient $r_{sp}$ was replaced by the upper (a) and the lower (b) endpoint of its 95% confidence interval. The relative importance was computed for each tree from the logistic regression model. The relative importance of the most influencing variable had a value of one and that of the other variables was scaled accordingly. For each variable, the relative importance values were aggregated by 1° latitude resolution and the points of the graph correspond to the average values. The grey areas around the curve correspond to the 95% confidence intervals. The acronyms MED., TEMP. and BOR. in grey bars refer to the Mediterranean, temperate and boreal biome, respectively. The white section corresponds to missing data.

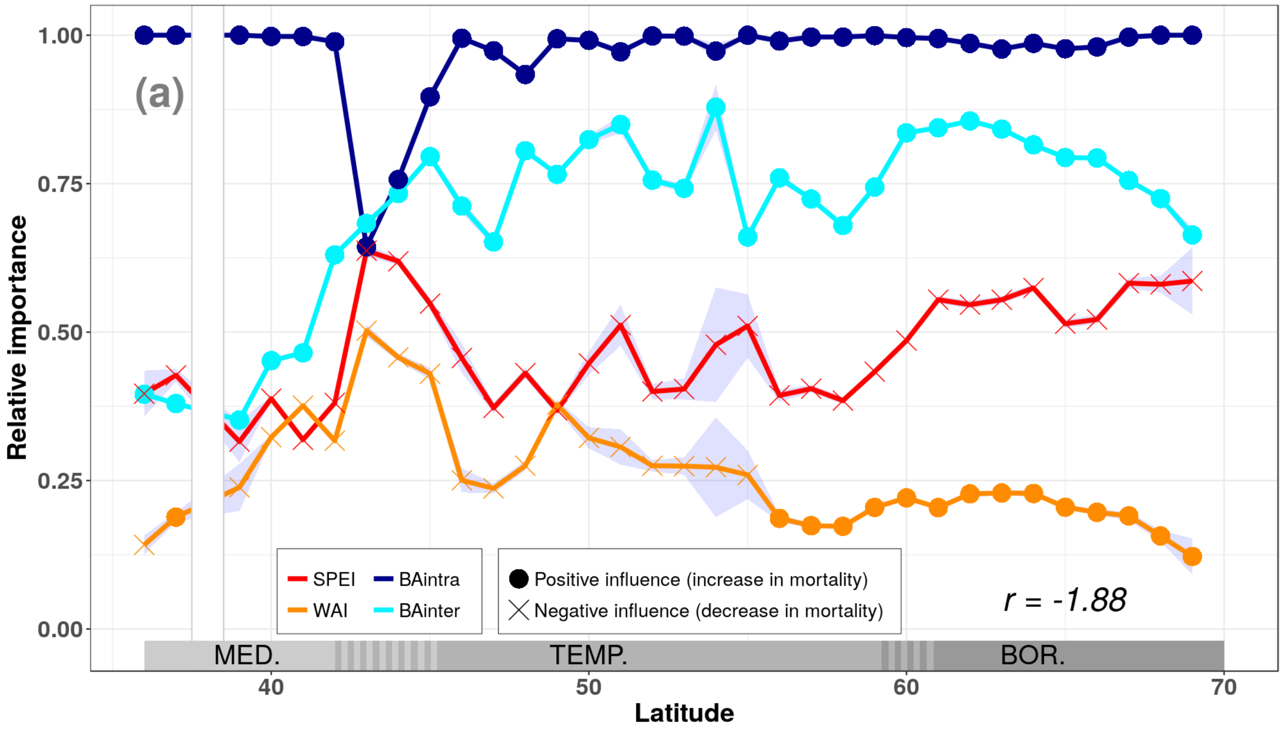


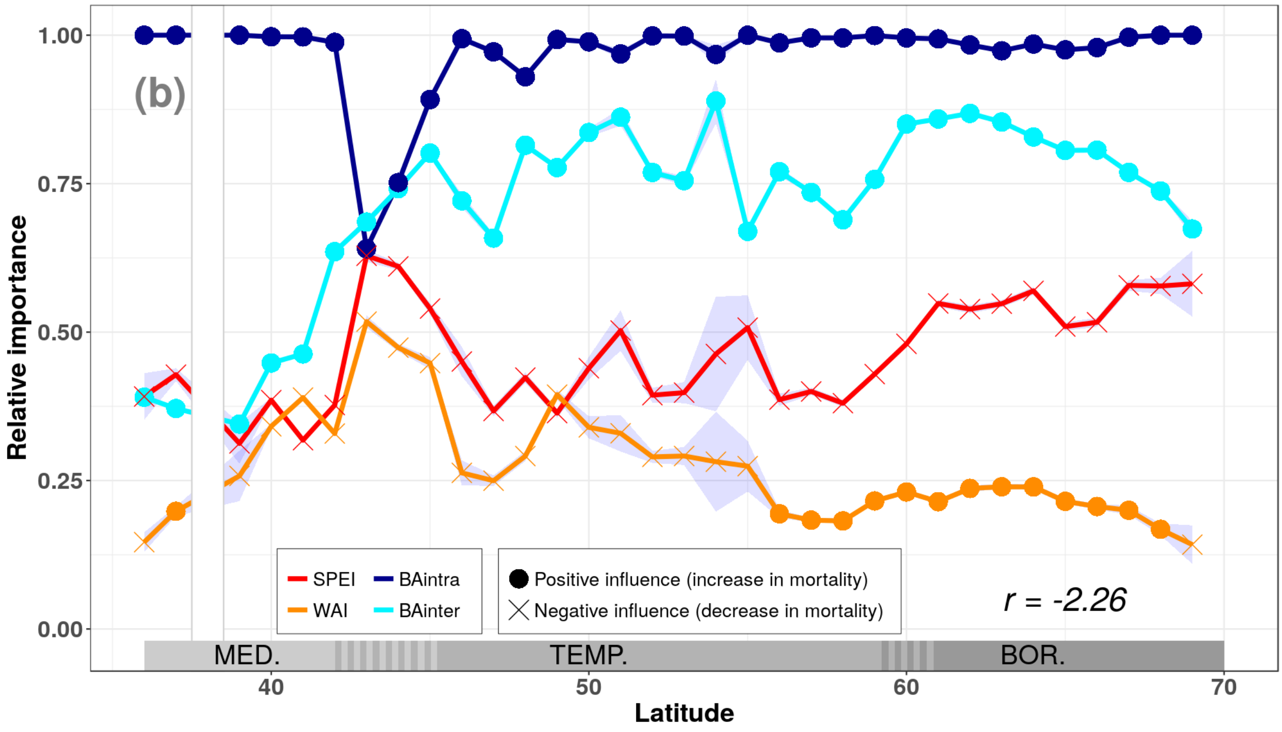

**Table S3.1 Coefficients estimates, standard errors and z values of species-specific logistic regression models.** A variable was considered significant if its z value was lower than -2 or higher than 2. $DBHnl_{sp}$ = $DBH$ × $r_{sp}$ log($DBH$) with $DBH$ = diameter at breast height (mm), *BAintra* = basal area of conspecifics ($m^2$ $ha^{-1}$), *BAinter* = basal area of heterospecifics ($m^2$ $ha^{-1}$), *SPEI* = Standardized precipitation evapotranspiration index (adimensional), *WAI* = water availability index (adimensional).

| | | ***P. sylvestris*** | | | | ***F. sylvatica*** | | | |
|---|---|---|---|---|---|---|---|---|---|
| | **Transformed variables** | **Model coefficient** | **Estimate** | **Std error** | **z value** | **Model coefficient** | **Estimate** | **Std error** | **z value** |
| **Biotic variables** | $\boldsymbol{DBHnl_{sp}}$ | $\beta_{3f}$ | 0.504 | 0.012 | 42.857 | $\beta_{3p}$ | 1.010 | 0.042 | 23.777 |
| | ***BAintra*** | $\beta_{4f}$ | 0.425 | 0.013 | 33.068 | $\beta_{4p}$ | 0.198 | 0.029 | 6.755 |
| | **log(*BAinter*)** | $\beta_{5f}$ | 0.292 | 0.013 | 21.705 | $\beta_{5p}$ | -0.167 | 0.034 | -4.888 |
| **Climatic variables** | **log(*WAI*)** | $\beta_{1f}$ | -0.064 | 0.017 | -3.712 | $\beta_{1p}$ | -0.225 | 0.038 | -5.856 |
| | **log(*SPEI*)** | $\beta_{2f}$ | -0.152 | 0.014 | -10.767 | $\beta_{2p}$ | -0.217 | 0.033 | -6.501 |
| **Interaction terms** | $\boldsymbol{DBHnl_{sp}}$ **× log(*WAI*)** | $\gamma_{1f}$ | 0.075 | 0.011 | 6.812 | $\gamma_{1p}$ | 0.262 | 0.042 | 6.263 |
| | $\boldsymbol{DBHnl_{sp}}$ **× log(*SPEI*)** | $\gamma_{2f}$ | 0.030 | 0.009 | 3.480 | $\gamma_{2p}$ | -0.097 | 0.037 | -2.642 |
| | ***BAintra*** **× log(*WAI*)** | $\gamma_{3f}$ | -0.107 | 0.012 | -9.235 | $\gamma_{3p}$ | -0.034 | 0.030 | -1.159 |
| | ***BAintra*** **× log(*SPEI*)** | $\gamma_{4f}$ | 0.164 | 0.011 | 14.657 | $\gamma_{4p}$ | 0.001 | 0.025 | 0.034 |
| | **log(*BAinter*) × log(*WAI*)** | $\gamma_{5f}$ | 0.031 | 0.012 | 2.563 | $\gamma_{5p}$ | 0.136 | 0.031 | 4.421 |
| | **log(*BAinter*) × log(*SPEI*)** | $\gamma_{6f}$ | 0.049 | 0.010 | 4.913 | $\gamma_{6p}$ | -0.036 | 0.028 | -1.292 |

**Figure S3.3 Histogram of the residuals and scatterplot of residuals versus fitted values for (a,b) Scots pine's model; (c,d) common beech's model.** (b,d) In binned residuals plots, $x$ fitted values are grouped in $n$ class with $n = \text{floor}(\sqrt{x})$ and average residuals are plotted against average fitted values in each group (Gelman & Hill, 2007). The grey lines indicate the 95% confidence interval within which binned residuals should be found. Plots were drawn using "arm" library (Gelman & Su, 2016) in R 3.3.3 (R Core Team 2017).

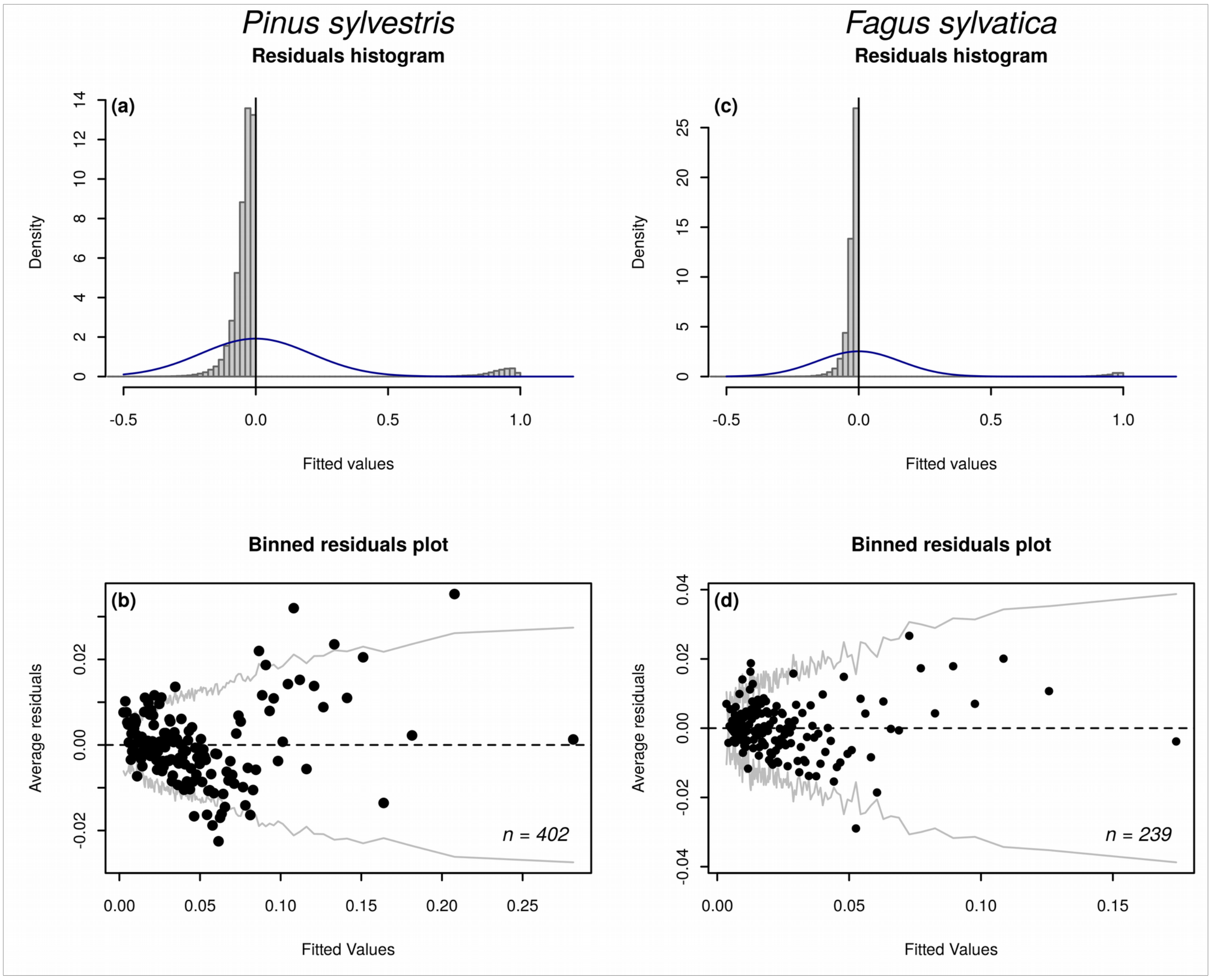

**Figure S3.4 Partial binned residual plots for (a) Scots pine's model and (b) common beech's model.** For each variable *x* observed variable values are grouped in *n* class with $n$ = floor($\sqrt{x}$) and average residuals are plotted against average observed variable values in each group (Gelman & Hill, 2007). The grey lines indicate the 95% confidence interval within which binned residuals should be found. Plots were drawn using arm library (Gelman & Su, 2016) in R 3.3.3 (R Core Team 2017).

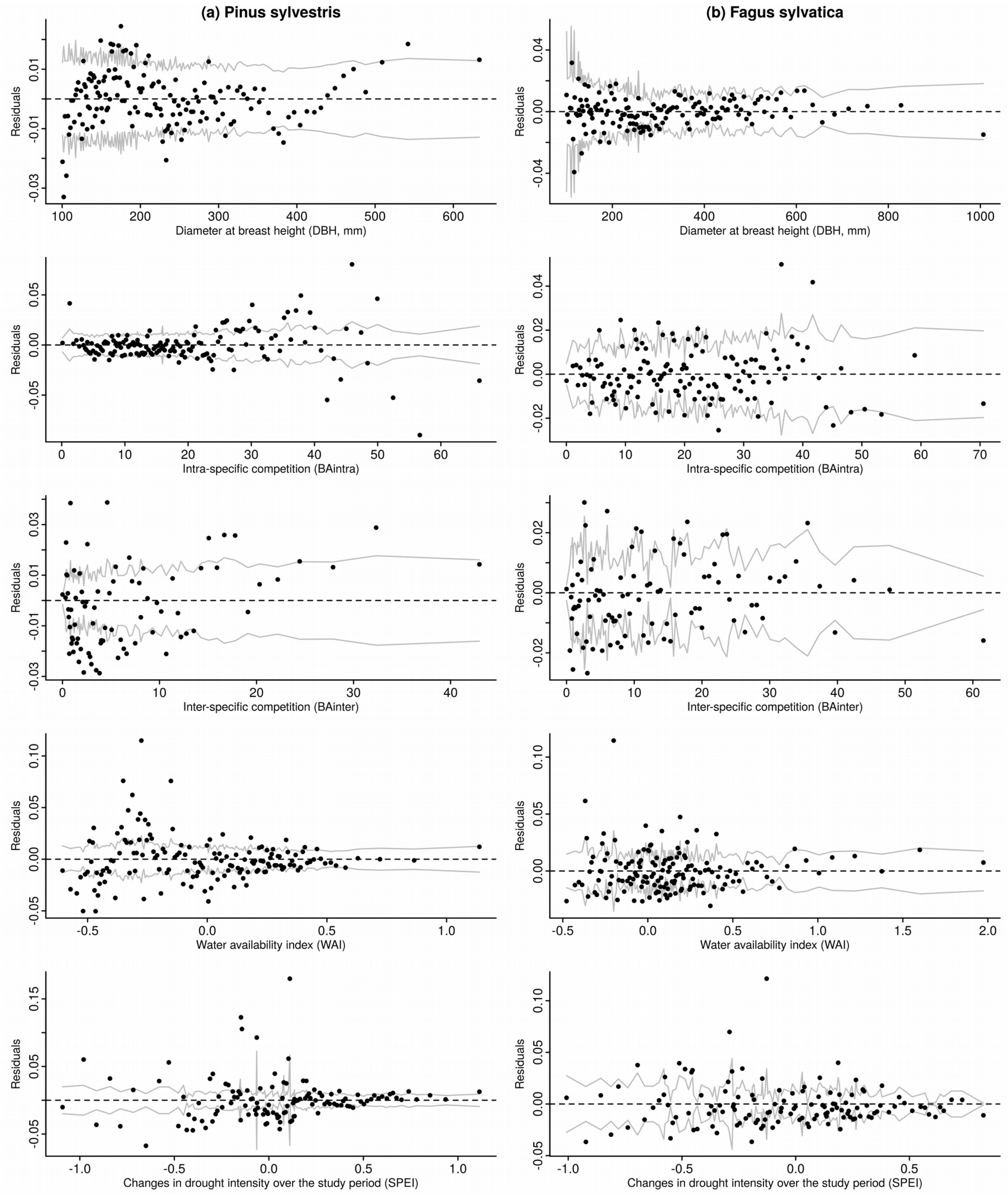

### Appendix S4. Interactions between basal area and drought-related variables not shown in the main figures for Scots pine.

**Figure S4. Interactions between conspecific basal area (i.e. *BAintra*), heterospecific basal area (i.e. *BAinter*), climatic drought intensity (i.e. *WAI*) and changes in climatic drought intensity over the study period (i.e. *SPEI*) on Scots pine probability of mortality.** These interactions were considered significant as their z value was lower than -2 or higher than 2 (see Table S3.1) but their effect was weaker than that of the interaction between *BAintra* and *WAI* (Fig. 2b). Scots pine mortality was predicted at three different levels of a given basal area (*BAinter* or *BAintra*) along a gradient of a given drought-related variable (*WAI* or *SPEI*) while the other predictors were fixed at their mean value. The three different levels of basal area correspond to their mean value, their 99.5$^{th}$ percentile and their 0.005$^{th}$ percentile (i.e. proxies of average, high and low competition, respectively). Caution has to be taken because the y-axis scale used in this Figure is not the same as that in Fig. 2 where the interaction was much stronger.

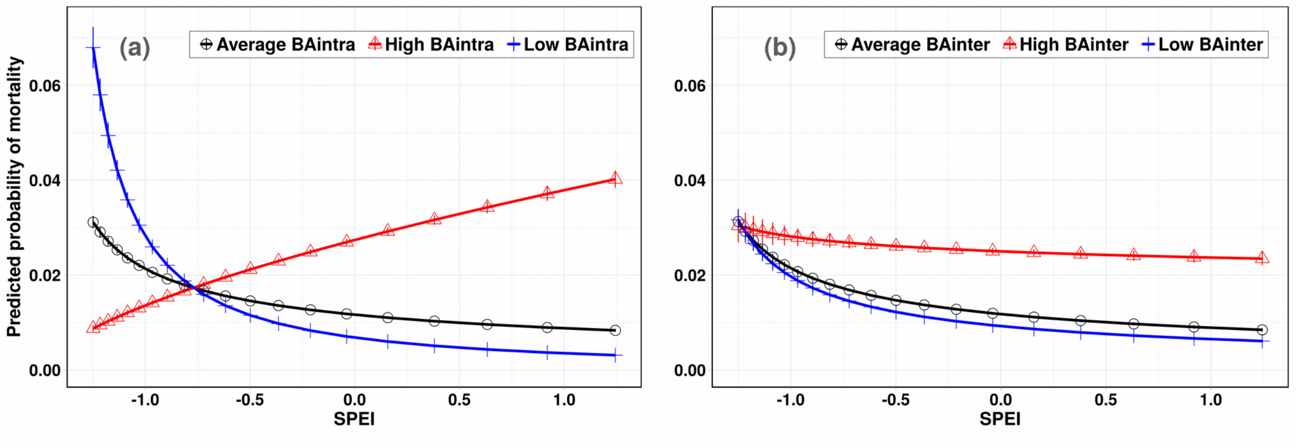


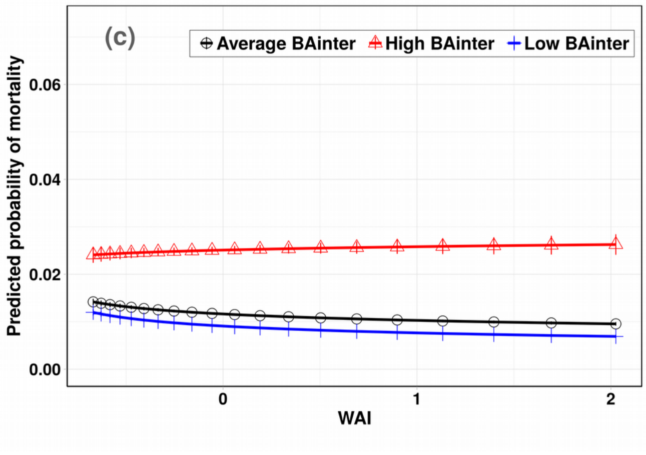
